# Date palm acclimates to aridity by diverting organic osmolytes for root osmotic adjustment in parallel with leaf membrane remodeling and ROS scavenging

**DOI:** 10.1101/2024.06.07.597900

**Authors:** Bastian L. Franzisky, Heike M. Mueller, Baoguo Du, Thomas Lux, Philip J. White, Sebastien Christian Carpentier, Jana Barbro Winkler, Joerg-Peter Schnitzler, Jörg Kudla, Jaakko Kangasjärvi, Michael Reichelt, Axel Mithöfer, Klaus F.X. Mayer, Heinz Rennenberg, Peter Ache, Rainer Hedrich, Maxim Messerer, Christoph-Martin Geilfus

## Abstract

**Highlight statement:** Osmotic strength of date palm roots increases with soil desiccation, for which the accumulation of organic osmolytes, such as sugars, is essential in complement to energetically cheap mineral osmotics.

Date palm (*Phoenix dactylifera* L.) is an important crop in arid regions that is well-adapted to desert ecosystems. To understand the remarkable ability to grow and yield in water-limited environments, experiments were conducted in a simulated desert environment with water-withholding for up to four weeks. In response to drought, root, rather than leaf, osmotic strength increased, with sugars contributing more to the osmolyte increase than minerals. Consistently, carbon and amino acid metabolism was acclimated toward biosynthesis at both the transcriptional and translational levels. In leaves, a remodeling of membrane systems was observed, suggesting changes in thylakoid lipid composition, which together with the restructuring of the photosynthetic apparatus, indicated an acclimation preventing oxidative damage. Thus, xerophilic date palm avoids oxidative damage under drought by combined prevention and rapid detoxification of oxygen radicals. Although minerals were expected to serve as cheap key osmotics, date palm also relies on organic osmolytes for osmotic adjustment of the roots during desiccation. The diversion of these resources away from growth is consistent with date palm’s strategy of generally slow growth in harsh environments and clearly indicates a trade-off between growth and stress-related physiological responses.

## 1. Introduction

Date palm (*Phoenix dactylifera* L.) is one of the most important crops in the semi-to hyperarid Arabian Peninsula. This is because it grows under a variety of extreme environmental conditions such as high soil salinity (Müller et al. 2023), heat and water scarcity (Arab et al., 2016; Du et al., 2019; Kruse et al., 2019; Du et al., 2023). Climate change will lead to increased global temperatures and reduced regional precipitation and, thus, to an increased frequency of extreme events such as heat waves and summer drought (Amin et al., 2016; Tabari and Willems, 2018; Saharwardi et al., 2023). Understanding drought effects and development of drought-tolerant genotypes is central to deliver increased crop yield in environments of limited water availability (Chaves und Davies 2010). However, plant responses to water deficit are complex (Puértolas et al., 2017) and dependent on specific drought scenarios, differing in the extent of water scarcity, irrigation placement (Dodd et al., 2008) and frequency (Boyle et al. 2016).

In response to drought, many plants undergo rapid physiological changes due to osmotic stress, including the increase in abscisic acid (ABA) as a prominent feature that might indicate the water use strategy of a species or genotype (Sreenivasulu et al., 2012). In addition to physiological functions in plant development, such as seed development, dormancy and storage of proteins and lipids (Finkelstein et al. 2002), ABA signaling mediates rapid decrease of stomatal aperture to reduced transpirational water loss. In turn, this decreased stomatal conductance causes a drop in leaf internal carbon dioxide (CO_2_) that negatively affects photosynthetic carbon fixation under water deficit (Cornic and Briantais, 1991; Brunner et al., 2015). Date palm is a relatively slow-growing crop with sclerophyllous leaves that exhibit comparably low stomatal conductance even under well-watered conditions (Kruse et al. 2019; Medrano et al. 2002). This conservative water use strategy (Kruse et al., 2019) is considered an evolutionary adaption to xeric environments (Mäkelä 1996). In addition to stomatal regulation, ABA triggers the expression of several stress-responsive genes in numerous plant species, involved in the accumulation of protective proteins such as dehydrins and other late embryogenesis abundant proteins (Ingram and Bartels, 1996; Verslues et al., 2006), and compatible osmolytes that contribute to lower the osmotic potential of the cell as part of osmotic adjustment (OA) (Chen and Murata, 2011; Yaish, 2015; Slama et al., 2015). OA is an acclimation response to dehydration that helps to maintain turgor and protect specific cellular functions by stabilizing protein and membrane structures. It constitutes an important factor in plant survival and yield (Blum, 2017). Plants might accumulate as osmolytes both inorganic minerals such as potassium (K^+^), chloride (Cl^-^), nitrate (NO ^-^) and others (Shabala and Shabala, 2011) as well as compatible organic compounds such as non-reducing sugars, polyols and amino acids (Chen and Murata, 2011; Slama et al., 2015). In comparison to OA by minerals, the accumulation of organic solutes takes place at an energetically higher cost (Yeo, 1983; Munns et al., 2020). Still plants exposed to soil water deficit osmotically adjust to declining soil water potential predominantly using organic compounds (Munns et al. 2020). Compared to the use of mineral solutes, this diverts significant amounts of photosynthetically fixed carbon and assimilated nitrogen from maintenance and growth processes. The type of organic osmolyte synthesized also impacts on related energy costs. The synthesis of sugars such as mannitol and sorbitol is cheaper in terms of water and nitrogen requirements than the synthesis of nitrogenous proline or glycine betaine. Recently, the cost in terms of carbon assimilated during the day has been calculated for OA in plants exposed to salt stress (Fricke, 2020; Munns et al., 2020), which initially acts largely as hyperosmotic perturbation (Golldack et al. 2011) and, thus, also requires OA. The diversion of 22% to 66% of daily assimilated carbon for glucose-mediated OA synthesis in leaves and roots, respectively, highlights the high cost of using organic osmolytes for OA (Munns et al., 2020).

The use of mineral osmolytes, although energetically more economical, is limited by the fact that high mineral ion concentrations, except for K^+^, can interfere with metabolic reactions in all compartments of the cytoplasm (Munns et al. 2016; Shabala 2013). It is assumed that mineral ions are mainly used to adjust the osmotic pressure in the vacuole, while in the cytoplasm it is balanced by the accumulation of compatible organic osmolytes (Shabala, 2013). For example, the prominent stress-induced accumulation of the compatible osmolyte proline makes a comparably small quantitative contribution to OA, but is associated with a protective effect on cellular functions and organs (Shabala and Shabala, 2011). Even under salt stress, when many mineral osmolytes in the soil solution are available as cheap osmolytes, usually only half of the required additional osmolytes for OA is covered by mineral osmolytes (Munns et al., 2020), generating an expensive demand for the synthesis of compatible organic solutes. Although energetically costly, the capability to osmotically adjust positively correlates with yield (Blum, 2017). Halophytes, such as date palm, are intrinsically characterized by tolerance to high tissue concentrations of mineral osmolytes such as K^+^, Cl^-^ and Na^+^. Date palm is able to increase the intake of K^+^ under high salinity (Mueller et al., 2023), illustrating a remarkable capacity for adjusting its transportome to achieve selective K^+^ intake in the face of hyperosmotic conditions and high external Na^+^ concentrations. The preferential use of energetically favorable mineral osmolytes for OA is a trait considered to improve plant growth in future climatic conditions (Munns et al., 2016; Munns and Millar, 2023). These adjustments also pose challenges for date palm productivity (Yaish und Kumar 2015; Allbed et al. 2017), despite date palm’s remarkable ability to withstand extreme environmental conditions.

Date palms mitigate effects of heat and drought employing proficient protein expression response comprising heat shock proteins and boosting of anti-oxidant activity to avoid oxidative stress (Safronov et al., 2017). In addition, isoprene synthetase (Ghirardo et al. 2021) and, consequently, isoprene emission (Arab et al., 2016) are upregulated, which are suggested to act as anti-oxidative mechanism that stabilizes thylakoid membranes. Seasonal drought results in variable metabolic responses (Du et al. 2021), including a shift in nitrogen metabolism that potentially facilitates a temporal accumulation of nitrogenous organic solutes, particularly in roots. However, there is limited information on drought-related OA of date palm, specifically regarding the use of energetically favorable mineral osmolytes versus more expensive organic solutes. To address this gap, a large-scale experiment that simulated the summer desert climate of the Arabian Peninsula was conducted and osmotic and metabolic acclimation in date palm roots and leaves were analyzed.

It was hypothesized that drought-exposed date palm **i)** preferentially relies on energetically favorable mineral osmolytes for OA, thus adapting the transportome in favor of mineral uptake, and **ii)** that its metabolism is acclimated by activating the anti-oxidative system to counteract increased ROS formation. To address these hypotheses, we exposed date palm cv. Khalas to continuous drought and analyzed OA by relating mineral, sugar, and amino acid concentrations to changes in osmotic strength. To follow changes in mineral transporter expression and metabolic acclimation underlying organic solute synthesis and other drought acclimation responses, we performed transcriptome, proteome and metabolite analyses of roots and leaves to elucidate how the xerophilic date palm thrives in arid deserts.

## 2. Material and Methods

### 2.1 Plant material and growth conditions

About two years old seedlings of micro-propagated *Phoenix dactylifera*, cultivar Khalas were purchased from Date Palm Developments Ltd. (Somerset, U.K.). Seedlings were planted in 5-L pots filled with 70% quartz gravel (3 – 5 mm diameter, Quarzwerke GmbH, Frechen, Germany) and covered by 4 cm soil substrate (Floragard Vertriebs-GmbH, Oldenburg, Germany). For the drought stress experiment, plants were transferred into walk-in phytotrons at the Research Unit of Environmental Simulation (EUS; Helmholtz Center Munich, Neuherberg, Germany), to simulate climatic conditions of the date palm’s natural habitat (12 h:12 h, light:dark; 40:20 °C; approx. 5:30% humidity, 600 μmol photons m^-2^ s^-1^ at shoot height) as described elsewhere (Du et al. 2023). Each plant was automatically irrigated by 50 ml of tap water every 4 hours. After 2 weeks of acclimation, water deprivation treatment was initiated by withholding watering for 3 days and then supplying 50 ml deionized water per day. Soil water contents were measured using a ML2 ThetaProbe connected to a HH2 moisture meter (Delta-T, Cambridge, UK). Soil moisture throughout the experiment was kept at 21.4 ± 7.5% and 10.7 ± 5.2% for control and water deprivation, respectively. The youngest fully expanded leaf and the whole root from each treatment were harvested for analysis of ions, elements, metabolites, and hormones 3, 10 and 31 days after onset of the treatment. Samples from the last sampling day were additionally subjected to transcriptomic and proteomic analyses. Plant materials were cut into small pieces, homogenized in liquid nitrogen and stored at −80°C until further analyses.

### 2.2 Biomass and tissue hydration

The fresh weight of whole shoots and roots was weighed at harvest. The corresponding dry weights were calculated using the fresh to dry weight conversion factor determined for each individual. This was the quotient of the weights after and before lyophilization of the collected homogenized tissue samples. Tissue hydration was determined as described previously (Du et al., 2021) and was used for calculating tissue concentrations of individual osmolyte classes.

### 2.3 Transcriptome analysis

RNA library preparation and sequencing were done at the Biomedicum Functional Genomics Unit (FuGU, Helsinki, Finland). The total RNA input amounts used for ribo-depletion were 200 ng for leaf (Ribo-Zero rRNA Removal Kit (Plant Leaf), Illumina) and 1 µg for root samples (Ribo-Zero rRNA Removal Kit (Plant Seed/Root), Illumina).

Directional sequencing libraries were constructed using Illumina’s ScriptSeq RNA Library Prep Kit. For this purpose, both mRNA and long non-coding RNA were sequenced from the samples. The resulting libraries were multiplexed and sequenced on an Illumina NovaSeq S1 flow cell 300 cycle flow cell (2×151 bp paired-end reads).

The quality of the raw RNAseq reads was analyzed with FastQC (http://www.bioinformatics.babraham.ac.uk/projects/fastqc). The trimming step was performed with Trimmomatic (Bolger et al., 2014) using the parameters lllILLUMINACLIP:Illumina_PE_adapters.fasta:2:30:10:8:true LEADING:3 TRAILING:3 SLIDINGWINDOW:4:20 MINLEN:60lll. The reads were mapped to a Trinity *de novo* assembly (Haas et al. 2013) using Kallisto (Bray et al., 2016). Assembly, construction, transcript annotation, and calculation of differentially expressed genes (DEGs) using lllEdgeRlll (Robinson et al. 2010) was performed as described previously (Mueller et al., 2023). DEG fold changes are given in base 2 logarithmic scale. Filtering for low read counts resulted in 70,285 transcripts. RNAseq data were submitted to EMBL-EBI-Annotare (https://www.ebi.ac.uk/fg/annotare) under the ArrayExpress accession number E-MTAB-14123.

### 2.4 Proteome analysis and computational integration in transcriptomic data

Protein extractions were performed following the phenol-extraction/ammonium-acetate precipitation protocol described previously (Carpentier et al., 2005; Buts et al., 2014). In brief, after extraction, 20 µg of protein were digested with trypsin (Trypsin Protease, MS Grade ThermoScientific, Merelbeke, Belgium) and purified by Pierce C18 Spin Columns (ThermoScientific). The digested samples (0.5µg/5µL) were separated in an Ultimate 3000 (ThermoScientific) UPLC system and then analysed in an Orbitrap ELITE mass spectrometer (ThermoScientific) equipped with an Acclaim PepMap100 pre-column (Thermo Scientific, Merelbeke, Belgium) and a C18 PepMap RSLC (Thermo Scientific) using a linear gradient of buffer A and B (0.300 μL/min). Buffer A was composed of pure water containing 0.1% formic acid; buffer B was composed of pure water containing 0.08% formic acid and 80% acetonitrile. The Orbitrap ELITE mass spectrometer (ThermoScientific) was operated in positive ion mode with a nanospray voltage of 1.8 kV and a source temperature of 275°C. The instrument was operated in data-dependent acquisition mode with a survey MS scan at a resolution of 60,000 for the mass range of m/z 375-1500 for precursor ions, followed by MS/MS scans of the top twenty most intense peaks with +2, +3, +4, and + 5 charged ions. All data were acquired with Xcalibur 3.0.63.3 software (ThermoScientific). For protein quantification, the software Progenesis® (Nonlinear Dynamics) was applied using all peptides for quantification as described in (Soares et al. 2018). We applied MASCOT version 2.2.06 (Matrix Science) against the assembled transcriptome using the search parameters parent mass tolerance of 12 PPM, fragment tolerance of 0.2 Da, variable modification by oxidation M, Deamidation NQ, fixed modification by carbamidomethyl C, with up to two missed cleavages allowed for trypsin.

The proteomic data (4,304 features, **Tab. S2**) were integrated into the transcriptomic data (51,494 features, **Tab. S1**). By searching the spectra against the mRNA database, transcript and protein abundances were linked to their genes (van Wesemael et al., 2018). The genes identified in both omics analyses were filtered for drought-related changes at the protein level and then examined for overrepresentation in Gene Ontology (GO). Overrepresented features related to plastid, anti-oxidant activity, and organic osmolyte metabolism were visualized in Cytoscape (Ma et al. 2021; Shannon et al. 2003).

### 2.5 Element analysis

The element concentrations of plant material were determined following acid digestion by inductively coupled plasma mass spectrometry essentially as described previously (White et al., 2012). In brief, 50 mg dried samples were weighed and digested in closed vessels using a microwave digester (MARS Xpress; CEM Microwave Technology, Buckingham, UK). Samples were first digested with 3 ml concentrated HNO_3_before the addition of 1 ml of 30% H_2_O_2_ to complete digestion. Digested samples were diluted to 50 ml with sterile MilliQ water before element analyses. Total K, Ca, Mg, P, S, Na, Cl, Fe, Mn, Zn, Cu and Ni concentrations were determined on digested material by ICP-MS (Nexion 1000, PerkinElmer, Waltham, MA, USA). Blank digestions were performed to determine background concentrations of elements and a tomato leaf standard (Reference 1573a; National Institute of Standards and Technology, NIST, Gaithersburg, MD, USA) was used as an analytical control.

### 2.6 Biochemical analyses

Foliar H_2_O_2_ was extracted and assayed as previously described (Mueller et al., 2023). *In vitro* glutathione reductase (EC 1.8.1.7) and dehydroascorbate reductase (EC 1.8.5.1) activities in leaves were determined as reported by (Arab et al., 2016). Total and oxidized glutathione (GSH), cysteine, γ-glutamylcysteine, total and reduced ascorbate were extracted and determined as described previously (Du et al. 2019). Relative abundances of water soluble low-molecular-weight metabolites in leaves were analyzed by GC-MS. For this purpose, metabolites were extracted, derivatized and separated as previously reported (Du et al., 2019). Phytohormone extraction and related LC-MS analysis was performed as described elsewhere (Vadassery et al. 2012; Dávila-Lara et al. 2021).

Amino acid concentrations were analyzed with modifications as described by (Döring et al., 2022). In brief, 30 mg of freeze-dried powder was dissolved in 1 ml buffer (0.12 M Li-citrate, 100 nM norleucin, pH 2.2). For extraction, samples were placed in an ultrasonic bath (Super RK 1028, Bandolin, Germany) cooled to 20 °C and constantly sonicated for 15 minutes. After centrifugation for 15 minutes at 15,000 xg, the supernatant was filtered through a 0.45 μm nylon filter and transferred to vials for measurement. Chromatography of unbound amino acids was performed over two hours on an amino acid analyser S433 using a 4.6 × 150 mm LCA K 07/Li cation-exchange column (Sykam, Eresing, Germany). Automatic post-column ninhydrin derivatization was applied before primary and secondary amino acids were detected photometrically at 570 and 440 nm, respectively.

Soluble sugar and starch concentrations were determined using the modified anthrone colorimetric method as described elsewhere (Li et al. 2013). In brief, 50 mg of dried and ground sample were mixed with 5 ml of 80% ethanol and heated to 80 °C for 30 minutes. After centrifugation at 5,000 g for 5 minutes, the supernatant was collected. This extraction was repeated and supernatants combined for soluble sugar analysis, while the precipitates were used for starch determination. To analyze starch, 2 ml of distilled water were added to the precipitates and heated in boiling water for 15 minutes. After cooling, 2 ml of 9.2 M HClO_4_ were added to hydrolyze the starch for 15 minutes. Then, 4 ml of distilled water were added and supernatant was collected after centrifugation at 5,000 g for 10 minutes. Starch hydrolyzation was repeated and supernatants were combined for analysis. Both extracts were analyzed spectrophotometrically at 620 nm using glucose as a standard. Starch levels were calculated by multiplying glucose concentration by the conversion factor of 0.9 (Osaki et al., 1991).

### 2.7 Osmolarity analysis

Osmolarity was measured using a semi-micro osmometer (Knauer ML, Berlin, Germany). To prepare the sample, 150 µL of the soluble sugar extract (section 2.5) was dried at 45 °C to evaporate the ethanol. Next, 150 µL of distilled water was added, and the solution was mixed vigorously before measuring according to the manufacturer’s protocol.

### 2.8 Statistical analysis

Biological replicates: biomass: n=5; ionome: n=5-10; phytohormones: n=5-10; metabolome, anti-oxidants, soluble sugars: n=5; proteome: n=4; RNA-sequencing: n=3. Data presented are from a large-scale experiment comparing different environmental factors with one control group; the control data are similar to those presented elsewhere (Mueller et al. 2023; Du et al. 2023). Data processing, analyses of variance, models and post-hoc tests (Tukey’s) were performed using R (R Core Team, 2013). Data were analyzed using linear models. Homogenity of data distribution was checked using the Shapiro-Wilk algorithm of ‘stats’. Inhomogeneously distributed data were logarithmically transformed and the normal distribution of the model residuals was verified or, if inadequate, the data were tested using the Kruskal-Wallis algorithm. Data sets with unbalanced sample sizes were evaluated using the HSD.test algorithm of ‘agricolae’ (Mendiburu and Yaseen, 2019) with the argument for unbalanced enabled. A significance threshold of *P*≤.05. was applied. For osmolyte correlation analysis a missing tissue hydration value (root_control_day1) was median-imputed from remaining replicates. Data were plotted using lllggplot2lll (Wickham, 2016) and ‘pheatmap’ (Kolde, 2015).

## 3. Results

### 3.1 Drought leads to accumulation of abscisic acid and activation of anti-oxidative metabolism in both roots and leaves

After three days of drought exposure, abscisic acid (ABA) concentrations in roots were increased to 60 ng g^-1^ DW, reaching a peak of 105 ng g^-1^ DW after 10 days of drought, while well-watered controls remained at about 16 ng g^-1^ DW throughout the experiment (**Fig. 1a**). A similar ABA pattern was observed in leaves (**Fig. 1b**), but at higher concentrations than in roots. Well-watered controls showed concentrations ranging from 90 to 110 ng ABA g^-1^ DW. Under drought, ABA increased to 190 ng g^-1^ DW after 3 days, reaching a maximum of 330 ng g^-1^ DW after 10 days and 250 ng g^-1^ DW after 31 days. Similar to ABA, the bioactive derivate of jasmonic acid, jasmonoyl-isoleucine (JA-Ile), accumulated in roots under drought (**Fig. 1a**). After three to 31 days of drought, JA-Ile was increased to about 25 ng g^-1^ DW compared to the control of about 10 ng g^-1^ DW. In leaves, higher JA-Ile concentrations were only observed after three days of drought, but at generally lower concentrations with a maximum at about 2 ng g^-1^ DW (**Fig. 1b**). For salicylic acid (SA), there was no significant response to drought in root tissue within one time point. In leaf tissue, on the other hand, a drought induced increase in SA concentration was observed after three days, but no further change at 10 and 31 days of drought exposure.

**Figure 1:**
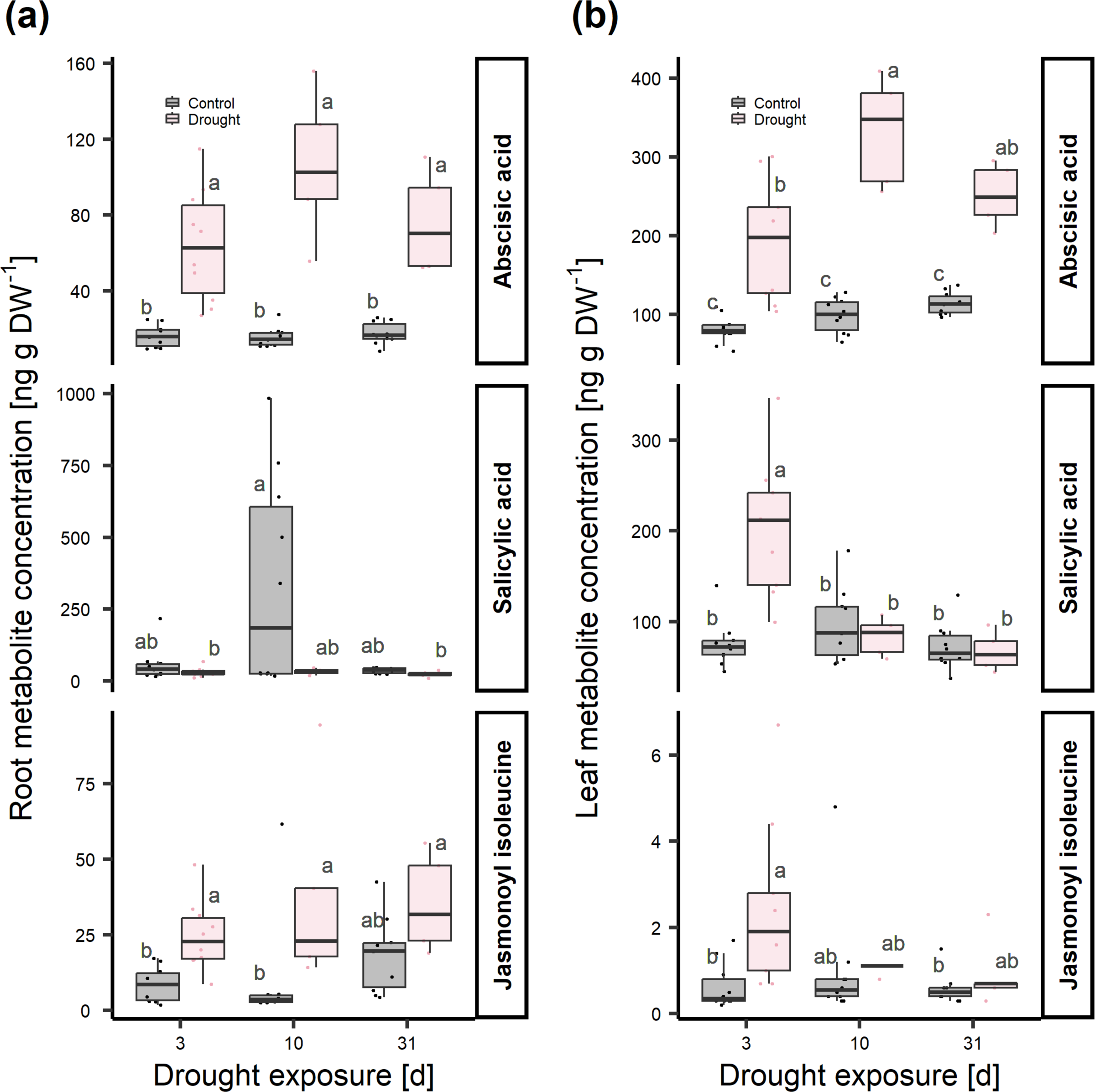
*Phoenix dactylifera* cv. Khalas accumulates abscisic acid in roots and leaves and jasmonoyl isoleucine mainly in roots under drought. Concentrations of abscisic acid and salicylic acid, and jasmonoyl isoleucine in (a) roots and (b) leaves of palms grown in phytototrons simulating a desert environment after 3, 10 and 31 days of well-irrigation or drought. Data presented as box plots featuring the maxima, 75 quartiles, medians, 25 quartiles and minima. Points shown represent raw data; n = 10-5; Tukey’s, *P* ≤ .05; different letters indicate significant differences of comparisons between water regime and timepoint.

Concurrent with the peak in ABA concentration after 10 days of drought exposure (**Fig. 2**), leaves showed an increase in H_2_O_2_ content that was still present after 31 days. With the increase in this ROS, metabolites and enzymes involved in ROS detoxification were upregulated, as evidenced by, e.g., increased foliar levels of ascorbic acid, glutathione, and glutathione disulfide at 10 and 31 days of drought exposure and increased glutathione reductase activity after 10 days. In roots, H_2_O_2_ did not accumulate at short or long drought exposure. Accordingly, the response of the anti-oxidative system was less pronounced in the roots than in the leaves. However, there were drought-related increases in the levels of dehydroascorbic acid, glutathione, and glutathione disulfide. Also, the activities of dehydroascorbic acid reductase and glutathione reductase were increased after three days of drought exposure, the glutathione reductase activity also after 10 days.

**Figure 2:**
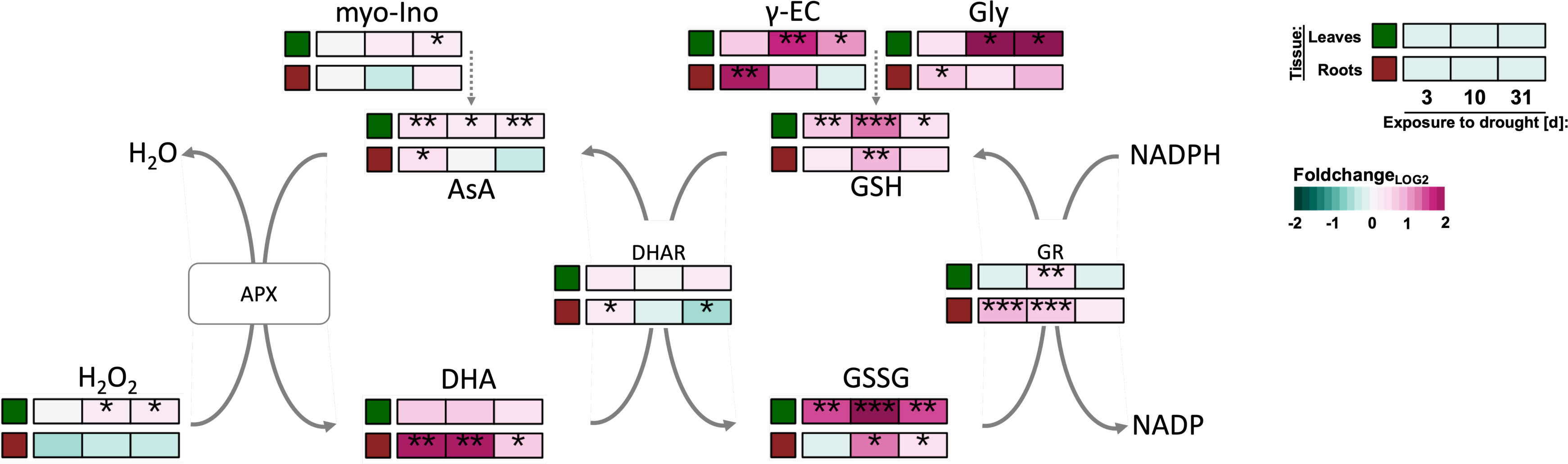
Anti-oxidative activity is stimulated in both roots and leaves of P*hoenix dactylifera* cv. Khalas under drought, yet foliar oxygen radicals accumulate. Response of metabolites and enzymes assigned to ascorbate-glutathione cycle in leaves and roots of date palms grown in phytototrons simulating a desert environment after 3, 10 and 31 days of well-irrigation or drought. APX, ascorbate peroxidase activity; AsA, reduced ascorbate; DHA, dehydroascorbate; DHAR, dehydroascorbic acid reductase activity; Gly, glycine; GR, glutathione reductase activity; GSH, total glutathione; GSSG, glutathione disulfide; myo-Ino, myo-inositol; y-EC, gamma-glutamylcystein. Means of the fold-change_LOG2_ are indicated by a color code (n = 5; t-test; *, *P* ≤ .05; **, *P* ≤ .01; ***, *P* ≤ .001).

The increased anti-oxidative demand as result of drought exposure was also evident at the protein level after 31 days of drought exposure. In roots, 10 proteins were annotated to the molecular function lllantioxidant activitylll (GO:0016209) (**Fig. 3a**), with ascorbate peroxidases 1, 2 and 3, NADPH-dependent thioredoxin reductase 2, glutathione S-transferases PHI9 consistently increased in response to 31 days of drought. Ascorbate peroxidase 1 and peroxidase superfamily proteins were shared with llllignin biosynthetic processlll (GO:0009809), which also showed increased levels of cinnamyl alcohol dehydrogenase 9, O-methyltransferase 1, aldolase-type TIM barrel family protein, and NAC domain transcriptional regulator superfamily protein. In leaves, catalase 2 was annotated to llloxidoreductase activitylll (GO:0016491), with two of the five detected homologs showing about 0.7-fold_LOG2_ decreased protein levels (**Fig. 3b**), while two homologous copper/zinc superoxide dismutases 2, annotated to the cellular component “plastid”, were about doubled after 31 days of drought.

**Figure 3:**
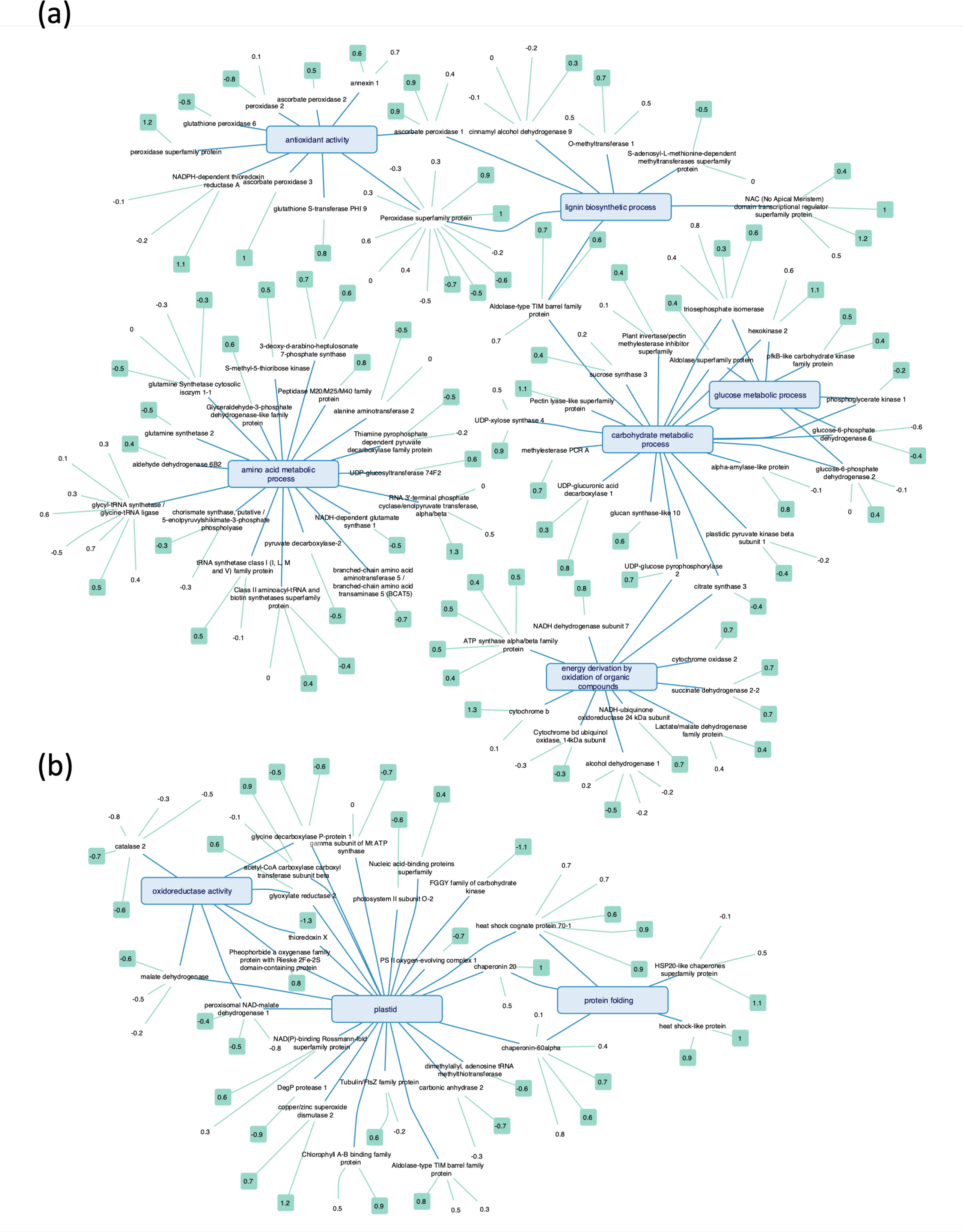
Changes in protein levels in roots and leaves of *Phoenix dactylifera* cv. Khalas in response to long-term drought exposure. Networks based on combined transcriptomic and proteomic data reveal strong responses of **(a)** roots and **(b)** leaves to 31 days of drought exposure. Transcriptome data were integrated with proteome data to match detected protein and underlying mRNA. Overrepresented Gene Ontology (GO) terms were identified based on drought-responsive proteins. GO terms with respect to the three domains lllMolecular Functionlll, lllBiological Processlll and lllCellular componentlll are shown in blue network hubs. Node labels on blue edges show individual gene annotations. Each of the rectangular-shaped boxes in the outer layer represent a protein feature annotated to the respective gene. Turquise coloring of the outermost node indicates a significant change at the protein level (n = 4; *P* ≤ .05), which could be either an increase or a decrease as indicated by a foldchange_LOG2_.

### 3.2 Drought inhibits plant growth and causes accumulation of osmolytes

There was no difference in root biomass between well-watered and drought-exposed date palms, with values of about 7 g DW per plant over the duration of the experiment (**Fig. 4a**). In contrast, a significant reduction in shoot mass was observed as a result of drought exposure, which was evident after 10 days and amounted to approximately 50% after 31 days compared to the shoot of well-watered control (**Fig. 4b**). The hydration of the root remained unaffected independent of drought exposure at about 3.25 g H_2_O g^-1^ DW (**Fig. 4c**). The leaves showed a lower hydration compared to the roots, with values around 1.8 g H O g^-1^ DW, which also remained unchanged despite the drought (**Fig. 4d**). In a first attempt to monitor the change in osmotic strength as a result of acclimation to drought exposure, total osmolytes were extracted and related to tissue water content. In the leaves, osmolyte concentrations were about 300 mM irrespective of the duration of drought exposure (**Fig. 4f**). Root osmolyte concentrations were lower and remained at about 125 mM under well-watered conditions (**Fig. 4e**). In response to drought exposure, the concentration increased over the course of drought exposure, resulting in a final concentration of approximately 275 mM after 31 days, representing a significant increase of 130% compared to control. To determine the types and individual contributions of osmolytes responsible for this significant increase in osmotic strength of date palm roots, we analyzed the concentrations of ions, soluble sugars and amino acids.

**Figure 4:**
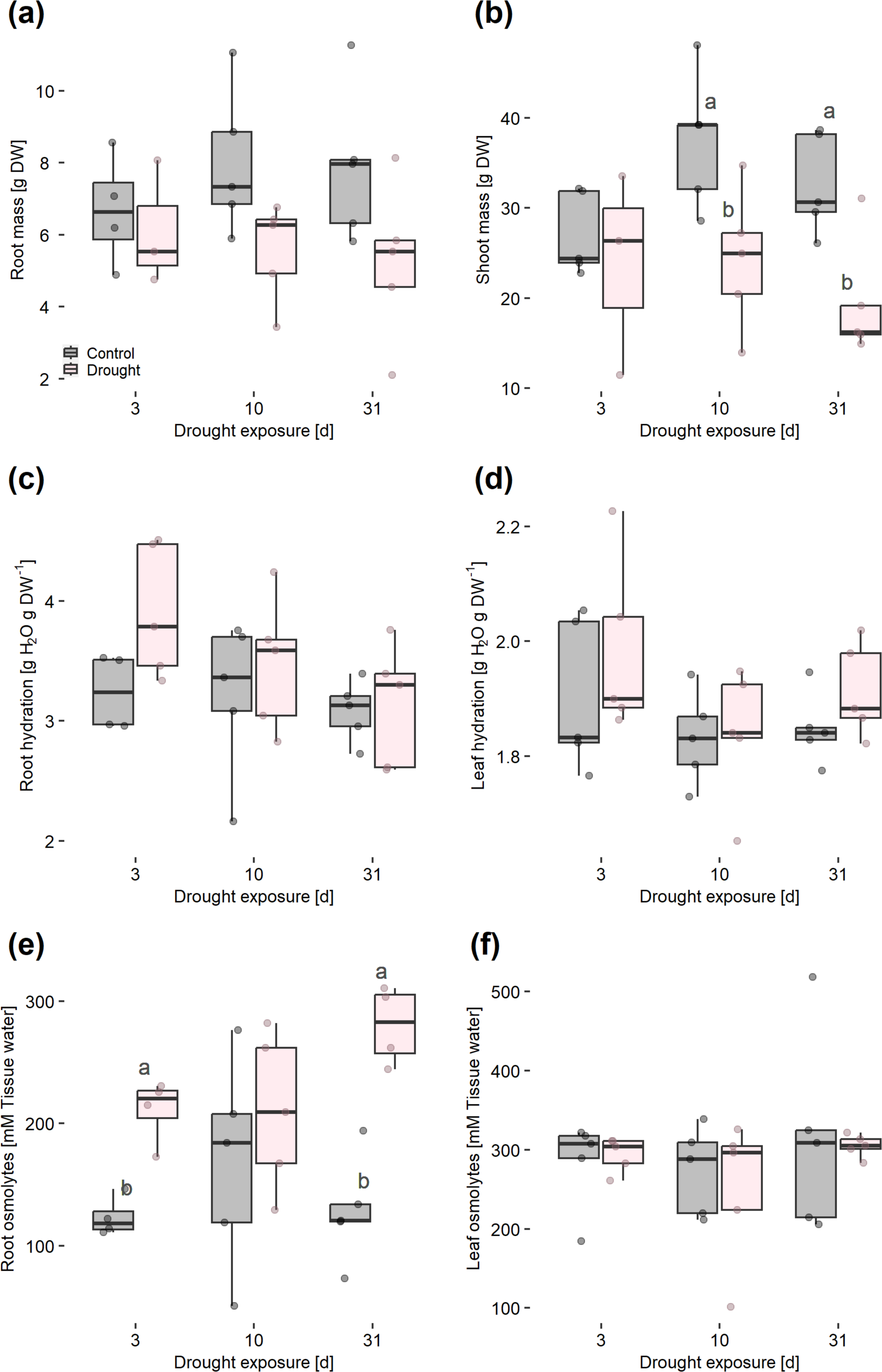
Root and leaf growth of *Phoenix dactylifera* cv. Khalas stagnates under drought stress, while tissue hydration remains unaffected with a parallel increase in osmolyte concentration. Biomass of **(a)** roots and **(b)** leaves, tissue hydration of **(c)** roots and **(d)** leaves, and osmolyte concentration of **(e)** of roots and **(f)** leaves of date palms grown in phytototrons simulating a desert environment after 3, 10 and 31 days of well-irrigation (control) or drought. Data presented as box plots featuring the maxima, 75 quartiles, medians, 25 quartiles and minima. Points shown represent raw data; n_BIOMASS_ = 5, n_Hydration_ = 5; n_Osmolytes_ = 5-4; Tukey’s, *P* ≤ .05; different letters indicate significant differences of water regime comparison within a timepoint; no letters, *P* > .05. Color code (a) for all panels.

### 3.3 Mineral osmolytes do not play a major role following drought exposure

Regarding halophytes, K^+^, Cl^-^ and Na^+^ are considered the major mineral osmolytes involved in cellular OA (Shabala and Shabala, 2011). The total concentration of these ions in roots was about 0.7 mmol g^-1^ DW without any significant changes in response the drought exposure (**Fig. 5a**). However, analysis of individual mineral concentrations in roots revealed that K^+^ concentrations increased by about 30% in roots after 31 days of drought compared to the well-watered control (**Fig. 5c**), while changes in Cl^-^ and Na^+^ concentrations were not significant (**Fig. 5e**). In leaves, the total mineral osmolyte concentration was about 0.8 mmol g^-1^ DW (**Fig. 5b**) and was not significantly affected by drought exposure as well. Leaf concentrations of K^+^ and Cl^-^ showed no significant changes (**Fig. 5d,f**), while Na^+^ concentrations differed after 10 and 31 days of drought exposure (**Fig. 5h**). However, the direction of change was not consistent and the relative contribution to the total mineral osmolyte concentration appeared negligible compared to K^+^ and Cl^-^. In both roots and leaves, the concentrations of other minerals such as Ca, Mg, P and S or Cu, Fe, Mn, Ni, Na and Zn showed no significant or marginal changes in response to drought exposure (**Fig. S1**).

**Figure 5:**
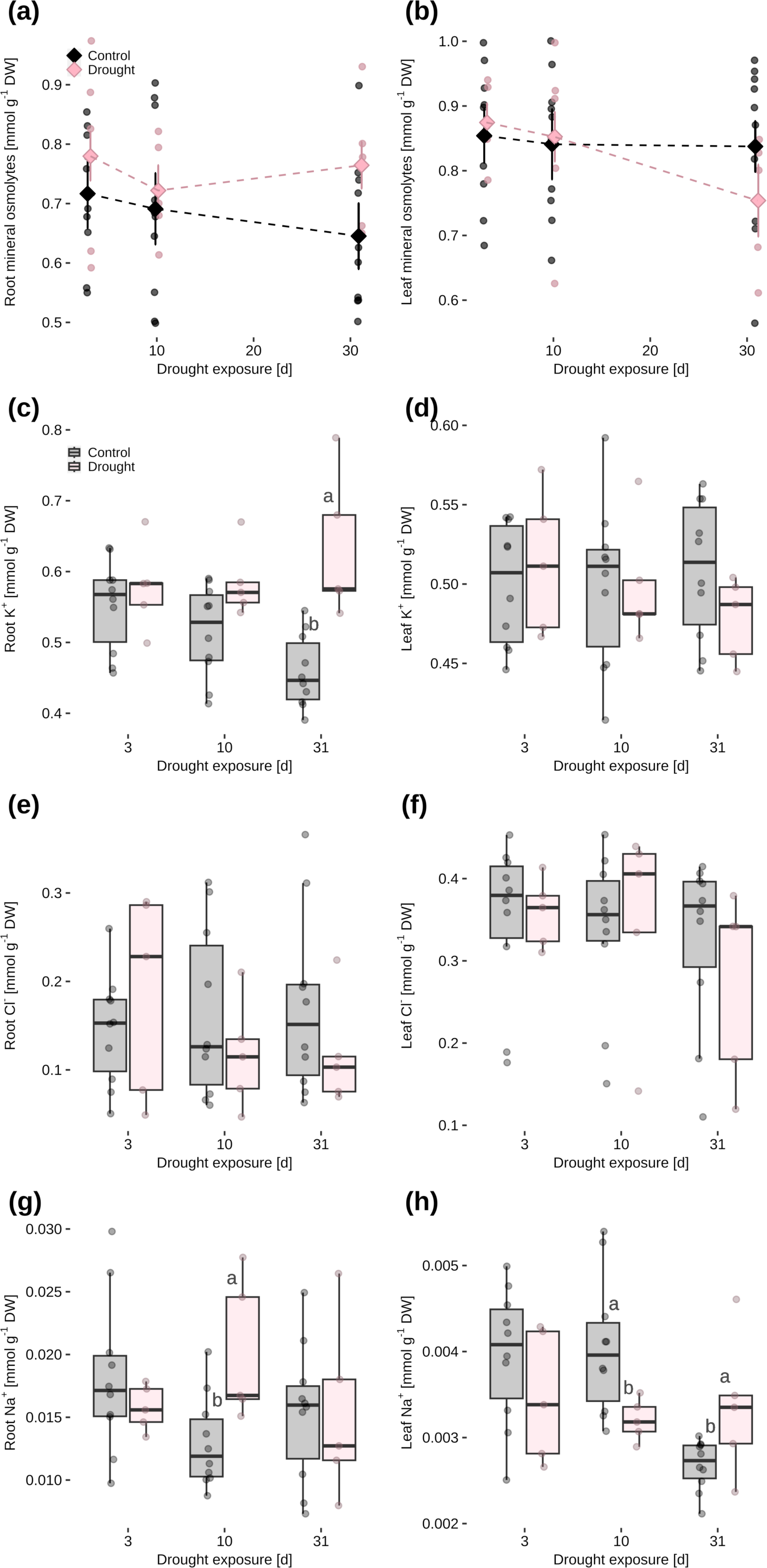
Mineral osmolyte concentration in roots of *Phoenix dactylifera* cv. Khalas remains unaffected during four weeks of drought exposure. Total concentration of the major mineral osmolytes of halophytes [potassium (K^+^), chloride (Cl^-^) and sodium (Na^+^)] in **(a)** root and **(b)** leaf and individual mineral concentrations in roots and leaves, respectively, of K^+^ **(c)** and **(d)**, of Cl^-^ **(e)** and **(f)**, and of Na^+^ **(g)** and **(h)** of date palms grown in phytototrons simulating a desert environment after 3, 10 and 31 days of well-irrigation (control) or drought. Data presented as box plots featuring the maxima, 75 quartiles, medians, 25 quartiles and minima. Points shown represent raw data; n = 10-5; Tukey’s, *P* ≤ .05; different letters indicate significant differences of water regime comparison within a timepoint; no letters, *P* > .05. Color code (a) or (c) for all panels.

To identify changes in mineral transporter expression associated with the increase in root K^+^ concentrations in response to drought, we analyzed the transcriptome of roots and leaves after 31-days drought exposure. Despite the drought-related increase in K^+^ concentration in date palm root, only few K^+^ transporter genes were differentially expressed after 31 days of drought. Among a total of 187 K^+^-transport related genes identified, only AKT2/3, a bidirectional channel, was found to be significantly up-regulated in the roots, but to a strong extent of about 40-fold (**Tab. S1a,b**). Consistent with the K^+^ concentrations in the leaves not affected by drought, 12 K^+^ transport related genes were ambiguously regulated, with 7 up-regulated (up to 14-fold_LOG2_) and 5 down-regulated (up to 3-fold_LOG2_).

### 3.4 Drought-related sugar accumulation in roots is favoured by upregulation of oligosaccharide biosynthesis

#### 3.4.1 Accumulation of oligosaccharides contributes to root osmotic adjustment

Measurement of total soluble sugar concentration in roots revealed a prominent increase in response to long-term drought exposure (**Fig. 6a**). From 3 to 10 days of drought, root sugar concentrations of about 0.4 mmol g^-1^ DW were not significantly higher than the control, but after 31 days of drought, there was a significant increase to about 0.5 mmol g^-1^ DW, a plus of about 60% compared to well-watered controls. The sugar concentrations in leaves were around 0.5 mmol g^-1^ DW and remained unaffected by drought exposure (**Fig. 6b**). Analysis of starch contents showed that leaves had higher starch levels than roots, with about 2% and 0.9% of DW, respectively (**Fig. 6c,d**). However, the drought treatment did not result in any significant changes in starch reserves for either leaves or roots, despite the latter showing a sharp increase in soluble sugars. To identify sugar compounds involved in the accumulation in roots after 31-days drought, we profiled the primary metabolites. Early changes in the sugar composition of roots were already apparent after three days of drought (**Fig. 6e**), with up to 0.8-fold_LOG2_ increases in galactinol and myo-inositol. After 10 days of drought, arabinose, gentiobiose, and trehalose were increased by 0.6 to 3-fold_LOG2_. Concurrent with the marked increase in total soluble sugars after 31 days of drought (**Fig. 6a**), a diverse range of sugar compounds was increased (**Fig. 6e**), with 2-fold_LOG2_ increases in fructofuranosyl-fructofuranose and gentobiose, and 1-fold_LOG2_ increases in idose, myo-inositol, and mannose, while the disaccharide sophorose was decreased. Although the leaves did not show an increase in total soluble sugars in response to drought exposure (**Fig. 6b**), the sugar composition was significantly altered (**Fig. 6e**). This change was characterized by 1 to 3-fold_LOG2_ increases in galactinol and raffinose, while arabinose was decreased, beginning three days after drought expsoure. Other sugars, such as fructose, 6-deoxy-mannopyranose, and glucose derivatives, also increased up to 1-fold_LOG2_, but not consistently.

**Figure 6:**
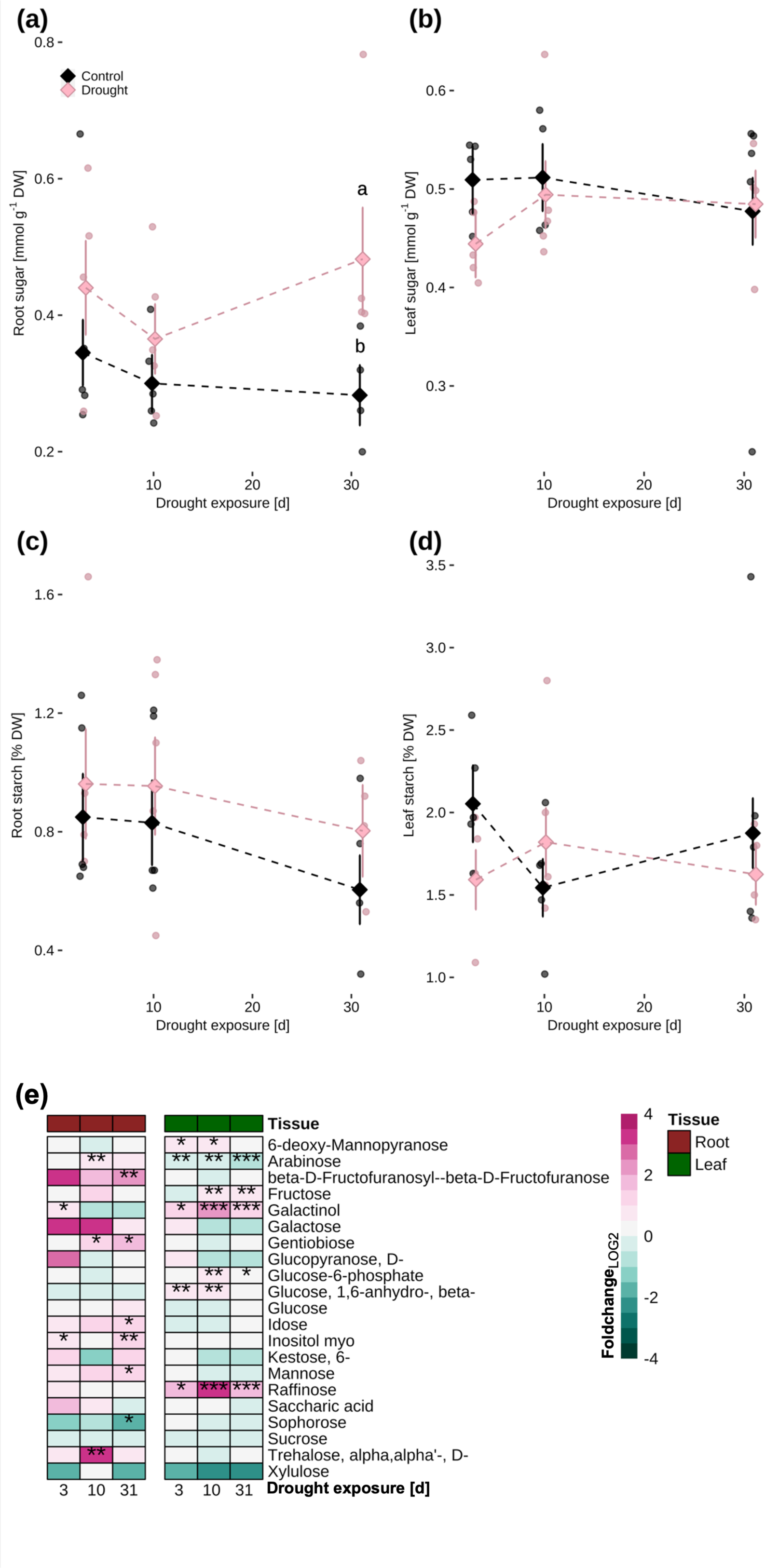
Sugar concentration increases in roots of *Phoenix dactylifera* cv. Khalas after four weeks of drought exposure while that of leaves remains unaffected. Concentrations in roots and leaves, respectively, of soluble sugars **(a)** and **(b)**, starch **(c)** and **(d)**, and **(e)** individual changes in sugar (alcohols and derivates) levels in roots and leaves of date palms grown in phytototrons simulating a desert environment after 3, 10 and 31 days of well-irrigation (control) or drought. Panel (a) and (b), means ± SE, points shown represent raw data; Tukey’s, *P* ≤ .05; different letters indicate significant differences of water regime comparison within a timepoint; no letters, *P* > .05. Panel (c), means of the foldchange_LOG2_ of the relative metabolite contents are indicated by a color code. Asterisks indicate the level of significance (*, *P*lll ≤ .05; **, *P* ≤ .01; ***, *P* ≤ .001); n = 5-4.

#### 3.4.2 Acclimation of root carbohydrate metabolism to drought favors oligosaccharide biosynthesis

In line with the marked increase in soluble sugar contents in roots (**Fig. 6a**), carbohydrate metabolism was significantly altered after 31 days of drought exposure. Transcripts of genes related to oligosaccharide metabolism showed a coherent increase of four galactinol synthases (3 to 7-fold_LOG2_) and one stachyose synthase (10-fold_LOG2_) (**Tab. S1f**) that both favor the synthesis of oligosaccharides (T Li et al., 2020). Consistently, analysis of proteomic responses revealed enrichment of 20 proteins that were annotated to the biological processes lllcarbohydrate metabolic processlll (GO:0005975) and lllglucose metabolic processlll (GO:0006006), with glucose-6-phosphate 1-dehydrogenase 2 and 6, hexokinase-2, phosphoglycerate kinase 1, aldolase superfamily protein and triosephosphate isomerase being shared between the two biological processes (**Fig. 3a**). Protein levels of glucose-6-phosphate dehydrogenase 2 and 6 homologs were ambiguously changed in response to drought, while other proteins of root carbohydrate metabolism, e.g. phosphoglycerate kinase 1, and citrate synthase 3 were significantly increased, except for plastidic pyruvat kinase beta subunit 1. Citrate synthase 3 and UDP-glucose pyrophosphorylase were shared with the term lllenergy derivation by oxidation of organic compoundslll (GO:0015980). Except of cytochrome bd ubiquinol oxidase 14 kDa subunit and alcohol dehydrogenase 1, the proteins related to this term were increased. Among these proteins were numerous homologs of ATP synthase aslpha/beta family protein (0.5-fold_LOG2_), cytochrome b (1.3-fold_LOG2_) and two homologs of succinate dehydrogenase 2-2 (0.7-fold_LOG2_).

In leaves, the transcript levels of genes related to carbohydrate metabolism and raffinose-family oligosaccharides such as galactinol synthases were ambiguously changed with one homolog 9-fold_LOG2_ decreased while two others were about 3 and 14-fold_LOG2_ increased (**Tab. S1f**), reflecting the increase in raffinose in leaves under drought (**Fig. 6c**). Genes related to starch metabolism were also differentially expressed. Transcripts of four genes involved in starch formation were coherently decreased, including two starch synthases with 8 and 2-fold_LOG2_ reduction. Some genes involved in starch degradation also had lower transcripts, e.g. phosphoglucan phosphatase and starch-debranching pullulanase. However, transcripts of two homologs of each of the alpha- and beta-amylases involved in starch degradation were increased by 2 to 5-fold_LOG2_, revealing an acclimation towards starch degradation in favor of oligosaccharides, although total starch content of the leaf remained unchanged (**Fig. 6d**).

### 3.5 Amino acid accumulation under drought is supported by enhanced biosynthesis

#### 3.5.1 Amino acids accumulate during long-term drought exposure

In both control and drought-stressed date palm roots, the total amino acid concentration was approximately 190 µmol g^-1^ DW after three and 10 days of exposure to drought (**Fig. 7a**). However, after 31 days of drought, the amino acid concentration was significantly higher by about 100 µmol g^-1^ DW, a plus of 50% compared to the control. In leaves of the well-watered control, the concentration of amino acids remained constant at around 20 µmol g^-1^ DW throughout the experiment (**Fig. 7b**). Under drought conditions, the concentration increased to a maximum of 45 µmol g^-1^ DW after 10 days, and then returned to a level similar to the control at approximately 30 µmol g^-1^ DW after 31 days. Subsequently, the contribution of individual amino acids to these changes was evaluated.

**Figure 7:**
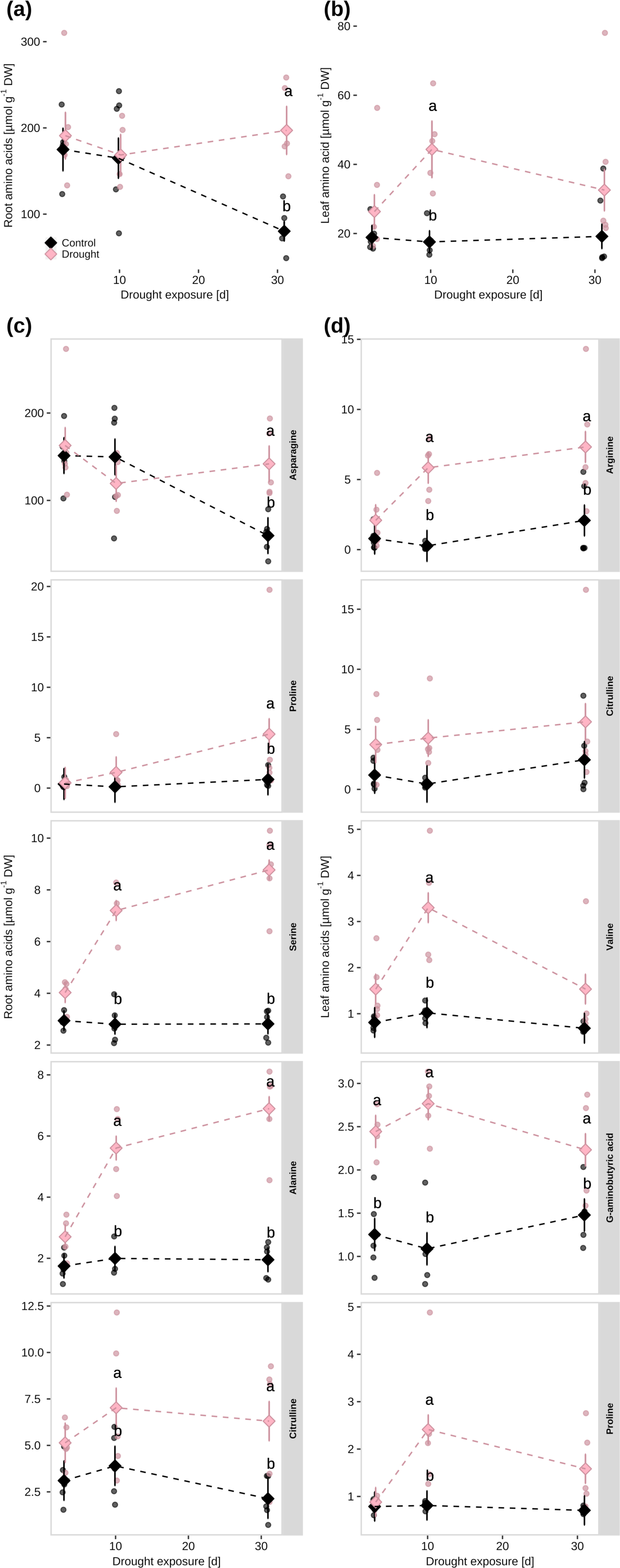
Amino acid concentration increases in roots of *Phoenix dactylifera* cv. Khalas already after 10 days of drought exposure. Total concentration of amino acids in **(a)** root and **(b)** leaf and top five individual amino acids with highest contributions to total amino acid accumulation under drought exposure in **(c)** roots and **(d)** leaves of date palms grown in phytototrons simulating a desert environment after 3, 10 and 31 days of well-irrigation (control) or drought. Means ± SE. Points shown represent raw data; n = 5; Tukey’s, *P* ≤ .05; different letters indicate significant differences in water regime comparisons within a timepoint; no letters, *P* > .05. Individual amino acids were selected according to the highest positive delta to the well-watered controls at the peak of accumulation (Fig. S3); full list of all amino acids detected provided in Fig. S2. Color code (a) for all panels to (d).

In roots, the amino acid composition was dominated by high concentrations of asparagine, with about 150 µmol g^-1^ DW (**Fig. 7c**). This concentration was similar to control conditions at 3 and 10 days of drought. However, after 31 days, the concentration declined in the control, but remained unaffected in drought-exposed roots, resulting in a plus of about 100 µmol g^-1^ DW. Other amino acids such as proline, serine, alanine, citrulline were among the amino acids detected, with concentrations ranging from 1 to 9 µmol g^-1^ DW. Increases related to drought exposure were observed, e.g., with a plus of about 5 µmol g^-1^ DW for proline, serine and alanine, while other amino acids showed lower or no drought-related increases. The amino acid composition of the leaves differed from that of the roots, mainly by a 10-times lower asparagine concentration than in the roots and was not affected by drought exposure (**Fig. 7d)**. Furthermore, exposure to drought led to an increase in the concentration of other amino acids in the leaves compared to the roots. These amino acids included arginine, citrulline, valine, gamma-aminobutyric acid and proline, which showed increases of up to 4 µmol g^-1^ DW. The concentrations of valine, proline, and gamma-aminobutyric acid increased transiently, reaching their peak after 10 days of drought. This pattern reflects the development of the total amino acid concentration, which peaked after 10 days of drought exposure with an 125 % increase compared to control (**Fig. 7b**).

#### 3.5.2 Acclimation of amino acid metabolism during long-term drought exposure

Drought-induced acclimation of amino acid metabolism was evident at the transcript and protein levels after 31 days of drought. In roots, three genes of amino acid metabolism were differentially expressed, with one down-regulated related to amino acid degradation, and two up-regulated related to the shikimate family and, thus, to the synthesis of aromatic amino acids (**Tab. S1g**). Drought-related changes in aromatic amino acid biosynthesis were also evident at the protein level (**Fig. 3a**). Among the 19 proteins annotated to lllamino acid metabolic processlll (GO:0006530), pyruvate decarboxylase 2 and 4, involved in aromatic amino acid catabolism, were coherently decreased (−0.5-fold_LOG2_). Other proteins involved in aromatic amino acid metabolism such as chorismite synthase (−0.3-fold_LOG2_), UDP-glucosyltransferase 74F2 (0.6-fold_LOG2_) and two homologous 3-deoxy-d-arabino-heptulosonate 7-phosphate synthases (0.6-fold_LOG2_) were ambiguously changed. In addition, protein levels of a glutamate synthase and two glutamine synthetases, catalyzing the conversion of glutamate to glutamine, were about halved after 31 days of drought. Proteins associated with protein biosynthesis, which play a role in the attachment of amino acids to their corresponding tRNAs, including glycyl-tRNA synthetase/glycine-tRNA ligase and tRNA synthetase class I family protein exhibited 0.5-fold_LOG2_ increases. In leaves, a total of 29 genes related to amino acid metabolism were differentially expressed (**Tab. S1g**), with 17 up-regulated, mostly related to biosynthesis, and 12 down-regulated related to both, biosynthesis and catabolism. No alterations in related proteins were observed following 31 days of drought exposure in comparison to the well-watered control.

### 3.6 Root osmotic adjustment involves parallel accumulation of sugars and amino acids, albeit in varying amounts

To assess the contribution of the individual osmolyte compounds to the OA observed in date palm roots in response to drought exposure (**Fig. 4e**), individual osmolyte concentrations were correlated with their sum, all related to the water content determined in the individual tissue fractions. In roots, both mineral and organic osmolytes increased with the total osmolyte concentration, with the highest levels observed after 31 days of drought exposure (**Fig. 8a,c,e**). Mineral osmolytes showed the highest significant correlation (*R* = 0.86, *P* < 0.001), followed by soluble sugars (*R* = 0.82, *P* < 0.001). The correlation between amino acids and the dependent variable was comparably lower (*R* = 0.42, *P* < 0.05). After 31 days, the roots of drought-exposed date palms exhibited a significant plus of 70 mmol soluble sugars per liter tissue water compared to well-watered controls. Thus, the contribution of sugars to the drought-related increase in root osmolytes was about 60%. Similarly, there was a plus of approximately 1.4 mmol amino acids per liter tissue water, while the increase in minerals was not significant. In leaves, similar positive correlations were observed between total osmolyte concentration and minerals and soluble sugars, respectively (**Fig. 8b,d**), but no effect of drought on leaf osmolyte concentration was evident (**Fig. 4f**; **Fig. 8b,d,f**).

**Figure 8:**
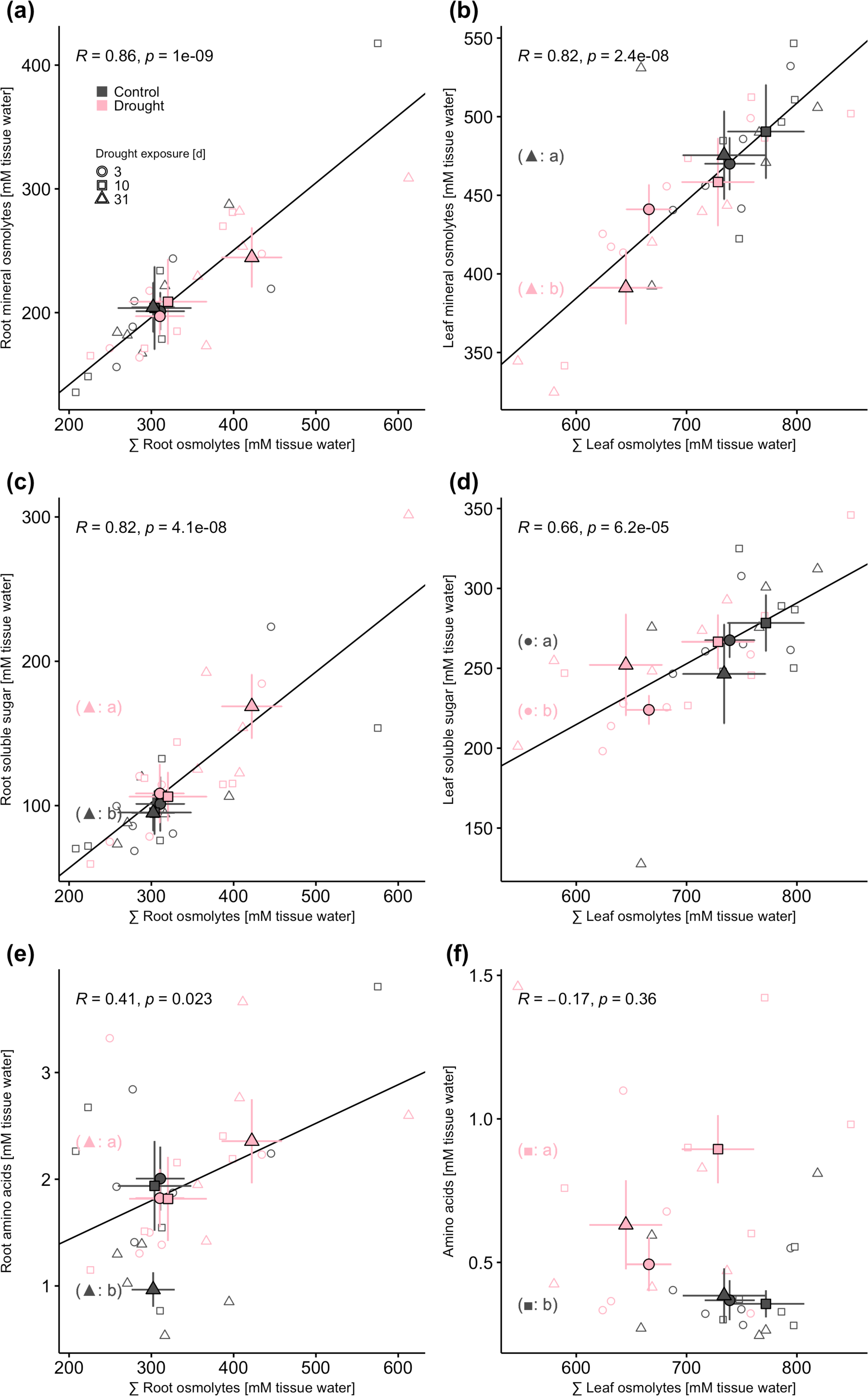
Total osmolyte concentration in roots of *Phoenix dactylifera* cv. Khalas increases with minerals, soluble sugars and amino acids. Correlation of the sum of measured osmolyte concentrations with concentrations of minerals **(a)** and **(b)**, sugars **(c)** and **(d)**, and amino acids **(e)** and **(f)** in roots and leaves, respectively, of date palms grown in phytototrons simulating a desert environment after 3, 10, and 31 days of drought. Measured osmolyte concentrations were related to cellular hydration of individual date palms. Pearson’s regression based on the raw data shown as open symbols; filled symbols represent mean ± SE; n = 5; Tukey’s, *P* ≤ .05; different letters in brackets indicate significant differences of individual osmolyte concentrations with respect to comparison of water regimes within a time point that is indicated by the shape code; no letters, *P* > .05. Shape and color codes (a) for all panels.

### 3.7 The photosynthetic apparatus and membrane systems are stabilized and restructured in leaves during drought acclimation

In response to 31-day drought exposure, the leaves showed a remarkable accumulation of numerous chaperones that were annotated to lllprotein foldinglll (GO:0006457). The heat shock-like protein and HSP20-like chaperones superfamily protein were doubled. Chaperonin 20 and 60alpha, and heat shock cognate protein 70 were shared with the cellular compartment lllplastidlll (GO:0009536; **Fig. 3**) and showed increased protein levels ranging from 0.5 to 1-fold_LOG2_. Among the 23 proteins annotated to lllplastidlll, photosystem II subunit O-2 and PSII oxygen-evolving complex 1 showed decreased abundance, while the abundance of chlorophyll A-B binding family protein was nearly doubled. With respect to plastid-associated lipids, drought-induced changes were evident at the transcript level (**Tab. S1c**). Transcripts of a set of eight genes encoding subunits of the lipid importer complex (TGD2,3,4), which facilitates the import of endoplasmic reticulum-derived lipids into the plastid (Fan et al., 2015), were coherently increased by about 2-fold_LOG2_, except for a TGD3 homolog, which was increased 8-fold_LOG2_. In addition, a 2-fold_LOG2_ increase in transcripts of two digalactosyldiacylglycerol synthase homologs and a 2-fold_LOG2_ up- and 6-fold_LOG2_ down-regulation of two homologous monogalactosyldiacylglycerol synthases suggested altered synthesis of thylakoid membrane-related galactolipids that are important for the stability of grana stacks.

Glycerolipids, such as phosphatidylcholine, a major constituent of lipid bilayers including thylakoid membranes, are synthesized via the Kennedy pathway. The transcript levels of phosphocholine phosphatase (−60-fold_LOG2_) and 1-acylglycerol-3-phosphate O-acyltransferase (AGAPT; −40-fold_LOG2_) were markedly decreased (**Tab. S1c**). Both are involved in the Kennedy pathway, where phosphocholine phosphatase hydrolyzes phosphorylated head groups of phosphocholine (Tannert et al., 2021) and AGAPT catalyzes the acylation of lysophosphatidic acid to phosphatidic acid, a critical step in synthesis of glycerolipids such as phosphatidylcholine and triacylglycerols (TAG). Other genes involved in glycerolipid biosynthesis were ambiguously regulated. These genes include glycerol-3-phosphate acyltransferases (GPAT1,9), involved in the synthesis of precursors for numerous glycerolipids, and three homologous diacylglycerol kinases (DAGK) that catalyze the phosphorylation of diacylglycerol to phosphatidic acid, which can trigger signaling pathways in response to environmental constrains (Kalachova et al. 2022) such as drought (Mane et al., 2007). Related to TAG synthesis, transcripts of three homologs of acyl-CoA:diacylglycerol acyltransferase (DGAT1,2) were 2-fold_LOG2_ up-regulated after 31 days of drought exposure. DGAT plays a key role in determining the flux of carbon into TAGs by catalyzing the only step exclusively dedicated to TAG synthesis (Jin et al. 2017; Liu et al. 2012), thus favoring TAG accumulation (Hatanaka 2022).

The remodeling of Kennedy pathway-derived glycerolipids in the Land’s cycle by means of deacylation-reacylation modifications is important for maintaining the integrity and functionality of cellular membranes. In this context, a marked −40-fold_LOG2_ reduction of transcripts encoding acyl-CoA:lysophosphatidylcholine acyltransferase (LPCAT) was evident. LPCAT modifies the fatty acid composition by catalyzing the re-acylation in the remodeling process. Conversely, lyso-phosphatidylethanolamine acyltransferase (LPEAT) was up-regulated approximately 2-fold_LOG2_. LPEAT is involved in an alternative acyl-editing pathway to the Lands’ cycle, which allows the exchange of fatty acids between phospholipids and the cytoplasmic pool of acyl-CoA (Klińska et al., 2020). Moreover, transcripts of three homologs of the UDP-glycose:sterol glucosyltransferase that are involved in the regulation of the sterol glycosylation level, were down-regulated 2 to 3-fold_LOG2_. Modification of sterols such as the glycosilation of membrane bound free sterols also affects membrane physical properties.

In addition to glycerolipids and sterols, sphingolipids are structural components in the plasma membrane and in endomembrane systems and contribute to maintaining their permeability and fluidity. Three genes involved in sphingolipid metabolism, *i.e.*, ceramide kinase, catalytic subunit 1 of serine C-palmitoyltransferase complex and inositol phosphosphorylceramide synthase, exhibited 2-fold_LOG2_ decreased transcript levels. Conversely, transcripts of ceramide glycosyltransferase, a key enzyme in sphingolipid metabolism that generates the precursor for all glycosphingolipids glucosylceramide, were about 4-fold_LOG2_ increased. Along with these changes, two homologous glycosylinositol phosphorylceramide mannosyltransferases, a Golgi-localized glycosylinositol phosphorylceramide-specific mannosyl-transferase (Ali et al. 2018), and a sphingolipid fatty acid 2-hydroxylase that catalyzes the 2-hydroxylation of the sphingolipid *N*-acyl chain (Hama, 2010), were up-regulated more than 2-fold_LOG2_, revealing that the overall expression of genes involved in the biosynthesis of sphingolipids was up-regulated in date palm leaves after 31 days of drought.

## 4. Discussion

### 4.1 Date palm mitigates drought-related radical generation by increasing anti-oxidant activity

Date palm is an important crop that is able to survive extreme environmental conditions such as heat and drought (Arab et al., 2016; Du et al., 2019; Kruse et al., 2019; Du et al., 2023). These conditions are prevalent in arid and semi-arid regions such as the Arabian Peninsula, where drought is a particular challenge (Almazroui et al., 2012; Almazroui et al., 2016) expected to worsen in future (Amin et al. 2016; Tabari und Willems 2018; Saharwardi et al. 2023). Already today, date palm productivity and fruit quality are challenged by this environmental constrain (Allbed et al., 2017; Ali-Dinar et al., 2023), emphasizing the need for a better understanding of date palm drought acclimation. To investigate the metabolic acclimation to drought with focus on OA, we analyzed date palms growing in simulated desert climate with a one-month period of drought. In response to these harsh environmental conditions, date palm shoot growth was significantly arrested (**Fig. 4b**), resembling previous observations of reduced leaf growth under field conditions (Ali-Dinar et al. 2023). The diminished shoot growth under drought is attributed to a reduction in the water potential of expanding leaf cells and leads to a higher root-to-shoot ratio (Davies, 2006), allowing the plant to allocate more resources toward root development to explore deeper soil layers for water uptake. However, no increase in root mass was observed after one-month of drought (**Fig. 4a**), indicating that the promoting effect on root growth was not yet pronounced.

Consistent with the observed shoot growth arrest, the drought-inducible stress hormone ABA, a negative regulator of plant growth (Yao and Finlayson, 2015) and stomatal opening (Müller et al., 2017), accumulated in both roots and leaves from the third day of drought exposure (**Fig. 1**). This pattern was also evident for JA-Ile, which is involved in stress acclimation (Riemann et al., 2015) and required for ABA biosynthesis in drought-exposed roots of Arabidopsis and citrus (Ollas et al. 2015; Ollas et al. 2013). In line, JA-Ile remained constantly elevated in drought-exposed date palm roots. In leaves, however, JA-Ile decreased after 10 days of drought exposure, despite increased ROS, which can stimulate JA signaling (Ismail et al., 2014). As a result of continuous stomatal closure and, thus, reduced leaf gas exchange and transpirational cooling, increasing leaf temperature and negative redox potential might cause generation of ROS (Rennenberg et al. 2006; Lee et al. 2012). This was evident in the leaves of date palm by increased levels of ROS (H_2_O_2_) (**Fig. 2**), as observed previously in date palms exposed to drought and heat (Arab et al., 2016; Du et al., 2019) or salt (Mueller et al. 2023). Consistently, anti-oxidant activity increased as evidenced by the increased abundance (**Fig. 3**) and activity of metabolites and enzymes involved in the Foyer-Halliwell-Asada cycle (**Fig. 2**) (Du et al., 2024), indicating a continued need to balance the cellular ROS homeostasis (Noctor et al., 2016) under drought.

### 4.2 Photosynthetic acclimation to drought involves stabilization and restructuration of antennae and lipid composition

Water deficit represents a significant challenge to photosynthesis, decreasing photosynthetic performance, while increasing the risk of ROS generation and oxidative damage to the photosynthetic apparatus (Pandey et al., 2023). Consistently, several changes in relation to the stabilization of the photosynthetic apparatus were evident in date palm leaves following one-month drought exposure, with prominent increases of chaperons (**Fig. 3b**). This response of date palm leaves to drought includes chaperonin 60 (Holland et al. 1998) and HSP70 (Aghaie and Tafreshi, 2020) that stabilize photosystems and can support chloroplast differentiation from plastids under elevated temperatures (Kim and An, 2013). Also constituents of photosystem antenna such as chlorophyll a/b binding proteins (CBPs) were increased. This increase might be regulated through ABA and JA (X-W Li et al., 2020), which were increased in date palm leaves following drought exposure (**Fig. 1**). CBPs play a role in plant acclimation to environmental cues (Ganeteg et al., 2004). In Arabidopsis, over-expression of a LHCB enhanced stomatal sensitivity to ABA (Xu et al. 2012), suggesting that the increase of CBPs in date palm leaves might be an acclimatory response to enhance guard cell ABA sensitivity to avoid transpirational water loss under continuous drought.

Moreover, remarkable remodeling of the lipid composition was evident in leaves after one-month drought exposure. During water deficit, which is often associated with elevated leaf temperature, membranes suffer oxidative damage. This makes it necessary to repair or replace lipids peroxidized by ROS (Benhiba et al., 2015; Anli et al., 2020; Shareef et al., 2020) to maintain stability and integrity of cellular membranes. Under drought, balancing of membrane fluidity and maintenance of protein interactions can be achieved by adjusting lipid composition in favor of unsaturated lipids, as seen previously in heat- and drought-exposed date palm leaves (Arab et al. 2016; Du et al. 2021; Safronov et al. 2017). Coherent increases in the expression of genes related to import of lipids into the plastids suggest increased demand for channeling endoplasmatic-derived lipid moieties for maintenance or replacement of thylakoid membranes under drought, a phenomenon observed under elevated temperature in numerous plant species (Li et al., 2015). In addition to disruption of membrane function and increased risk of lipid peroxidation, low cellular water disrupts grana stacking (Yu et al. 2021) and thus photosynthesis. Consistent with the pronounced thylakoid lipid remodeling observed in numerous other species in response to drought (Yu et al., 2021), date palm showed altered gene expression that might favor the increase in the ratio of di-galacto-to mono-galactolipids in thylakoid membrane (**Tab. S1c**), which is suggested to enhance stability in response to abiotic stress (Yu et al. 2021). The combined remodeling of thylakoid membranes and the restructuring of the photosynthetic apparatus might underlie the previously observed drought acclimation of date palm photosynthesis, *i.e.* the reduction in electron transport rate at reduced photosynthetic carbon fixation (Kruse et al., 2019), with reduced risk of photooxidative damage in hot, dry environments (Mäkelä 1996; Kruse et al. 2019).

Similar to the remodeling of thylakoid lipids, the composition of the plasma membrane might be modified to mitigate drought effects, e.g., by adjusting polar lipid head group proportions in favor of the non-bilayer forming phosphatidylethanolamine via the Kennedy pathway (Larsson et al., 2006). However, the observed changes in date palm expression of genes related to this pathway as well as those of the Land’s cycle, involved in glycerolipid remodeling, did not provide a clear picture in this regard. However, increased flux of carbon into TAG synthesis was suggested by increased expression of related enzymes after 31-day drought exposure. This response might aid in sequestering toxic lipid intermediates that accumulate due to membrane lipid remodeling (Lu et al. 2020). Date palm leaves also showed evidence for increased sphingolipid biosynthesis and glycosylation of lipid head groups in response to drought (**Tab. S1c**). An accumulation of glycosylinositolphosphoceramides and acylated steryl glycosides in drought-exposed plants has been associated with an increase in the content of sugar head groups in membranes, which helps to prevent protein precipitation upon desiccation (Tarazona et al., 2015). The combined accumulation of sterols and sphingolipids might be attributed to the formation of microdomain structures such as lipid rafts (Liu et al. 2021; Tapken und Murphy 2015). In addition to facilitating protein clustering and serving as a focal point for cellular signaling pathways, the temperature stability and activity of embedded protein complexes might be enhanced (Beck et al., 2007). Further analysis is needed to fully understand drought-related lipid remodeling in date palm.

### 4.3 Organic osmolytes make a major contribution to the osmotic adjustment of the roots

In addition to stomata closure, ABA accumulation also leads to various acclimatory responses involved in stress mitigation (Yoshida et al. 2014), such as OA (Shabala and Shabala, 2011). Because root cells contain more water per dry mass than leaves (**Fig. 4c,d**), OA of roots requires more osmolytes and, thus, is energetically more demanding (Munns et al., 2020). Accordingly, drought-related osmolyte accumulation was more pronounced in date palm roots than in leaves. In roots, a modest accumulation of the energetically cheap mineral osmolyte K^+^ was detected (Raven, 1985; Shabala and Shabala, 2011; Munns et al., 2020), which did not have a significant effect on OA (**Fig. 8a**). Consistent with the moderate K^+^ increase, only minor modifications to mineral transport of K^+^ were apparent (**Tab. S1a,b**), compared to the extensive adjustments observed under hyperosmotic saline conditions that enabled accumulation of K^+^ despite high Na^+^ and Cl^-^ concentrations in the soil solution (Mueller et al. 2023).

The use of organic compounds such as sugars and amino acids is more expensive than the use of mineral osmolytes because carbon skeletons have to be diverted from maintenance and growth processes (Shabala and Shabala, 2011; Munns et al., 2020) and, in the case of amino acids, represent a sink for often limited nitrogen resources (Raven 1985). Consistent with the known pattern that drought stress leads to an rapid increase in soluble sugars to minimize dehydration damage (Kaiser, 1987; Slama et al., 2015; Fàbregas and Fernie, 2019), date palm accumulated soluble sugars in the roots following drought exposure (Safronov et al., 2017; Anli et al., 2020), with the highest plus of 70 mM (**Fig. 8**) among the analyzed osmolytes. Consistently, roots showed marked up-regulation of galacatinol and stachyose synthases involved in oligosaccharide biosynthesis (Li et al. 2020a), while in the leaves, starch hydrolysis was up-regulated and starch formation down-regulated (Lloret et al., 2018; Ribeiro et al., 2022). Similar to previous observations (Du et al. 2021; Du et al. 2019), a shift in leaf sugar composition toward raffinose was apparent in the present study (**Fig. 6e**). In drought acclimation, raffinose accumulation might be related to membrane and protein stabilization by maintaining hydrophilic interactions (Fernandez et al., 2010) during water deficit conditions and acting as an anti-oxidant (ElSayed et al., 2014; Slama et al., 2015). Numerous sugars other than the typical free sugars, such as sucrose, glucose, and fructose (Fàbregas and Fernie, 2019), accumulated in roots (**Fig. 6e**), including gentiobiose, idose, myo-inositol, mannose, and the non-reducing disaccharide trehalose, a highly soluble and unreactive sugar that accumulates in desiccation-tolerant species during dehydration (Ilhan et al., 2015). Temporary increases in gentiobiose and trehalose were observed also in date palms exposed to hyperosmotic salt conditions. However, the total amount of sugars only slightly increased (Al Kharusi et al., 2019) or remained unaffected in date palm roots exposed to salinity (Mueller et al. 2023), similar to salt-exposed barley, where sugars also modestly contributed to OA (Annunziata et al., 2016). These findings suggest that the remarkable accumulation of soluble sugars with the underlying reprogramming of carbohydrate metabolism in the roots of date palm represents an acclimatory response specific to drought.

The contribution of amino acids to drought-related OA in roots (**Fig. 8e**) was significantly lower compared to the contribution of sugar osmolytes. To synthesize amino acids, nitrogen must be assimilated, which requires additional energy associated with an additional transpirational water loss of estimated 15 x 10^3^ mol H O (mol N)^-1^ for nitrate reduction (Raven, 1985), representing a critical investment under water scarcity. Although amino acids increased with osmotic strength in roots, the often accumulating compatible osmolyte proline (Slama et al., 2015) was not among the major contributors. Instead, the increase was mainly due to the generally high concentrations of asparagine that can serves as a storage compound for assimilated nitrogen due to its high nitrogen to carbon ratio and might accumulate, e.g., in response to stress-related reduction in protein biosynthesis (Lea et al., 2007). In date palm (Yaish, 2015; Safronov et al., 2017; Shareef et al., 2020; Du et al., 2021) and other halophytes (Slama et al., 2015), free amino acids accumulate under drought (Slama et al., 2015). This accumulation can be a consequence of numerous factors such as water deficit and ABA-related protein degradation (Huang und Jander 2017). However, concentrations of soluble proteins were previously not affected by drought (Du et al., 2023), suggesting no excessive drought-related protein degradation. Therefore, the observed stimulation of amino acid metabolism during drought strengthens the argument that also in date palm amino acids might act as a metabolic cache for organic nitrogen under stress (Hildebrandt, 2018).

### 4.4 Date palm does not exhibit a preference for cheap mineral osmolytes

Halophytes such as date palm are thought to tolerate higher cellular mineral concentrations (Flowers et al., 2015). Therefore, OA could be achieved primarily through the use of mineral osmolytes with low risk of ion toxicity. This trait might be exploited to achieve cost-effective OA through mineral uptake and sequestration (Shabala and Shabala, 2011; Munns et al., 2020), while reducing the need to synthesize organic compounds and divert them from growth. Date palm has been shown to remarkably control the uptake and long-distance transport of Na^+^ and Cl^-^ while promoting the uptake of K^+^ when growing in a hyperosmotic saline environment (Mueller et al. 2023). However, despite a marginal increase of K^+^ in roots, date palm showed no extensive mineral use for OA upon drought, while high amounts of sugars and much lower amounts of amino acids were diverted from maintenance and growth. This result confirms the primary use of organic osmolytes for drought-related OA (Munns et al., 2020) also in date palm, although increased uptake and sequestration of energetically less expensive mineral osmolytes would be a strategy preventing stress-induced growth arrest due to energy limitation (Shabala und Shabala 2011; Munns et al. 2020).

## 5. Conclusion

Date palm is a useful non-model crop species for studying tolerance to extreme environments. It is able to survive in arid deserts because it **(i)** restructures the photosynthetic apparatus to avoid formation of ROS, **(ii)** promotes anti-oxidative metabolism and **(iii)** osmotically adjusts the roots to low soil water potential. Energetically expensive organic osmolytes are essential for osmotic adjustment. Halophytes, with their expected higher intrinsic tissue tolerance, could achieve osmotic adjustment mainly by using mineral osmolytes with less risk of ion toxicity. Contrary to this assumption, date palm relies on sugars and amino acids rather than promoting mineral uptake. This strategy might bind resources to the detriment of growth and maintenance processes.

## Acknowledgements / Funding

P.W., K.F.X.M., J.Ku., J.Ka., H.R. and R.H. were supported by grants from the King Saud University, Riyadh, Saudi Arabia. M.M. was funded by the German Research Foundation (DFG) - TRR 356. C.-M.G. was funded by the German Research Foundation (DFG) - Projektnummer 499216091. In Memoriam of Professor Philip J. White (1960 - 2023) and Professor Jaakko Kangasjärvi (1960 – 2024).

## Author contributions

B.L.F., M.M., H.M.M., B.D., T.L., S.C.C., P.W., M.R., A.M., J.B.W., J.-P.S., C.-M.G. and P.A. performed the research and analyzed the data. J.-P.S., K.F.X.M., J.Ku., J.Ka., H.R., C.-M.G., P.A., and R.H. designed the study. B.L.F. wrote the draft of the manuscript with input from M.M., P.A., C.-M.G. and H.R., which was revised by all the authors.

## Abbreviations

- 1-acylglycerol-3-phosphate O-acyltransferase, AGAPT
- Abscisic acid, ABA
- acyl-CoA:lysophosphatidylcholine acyltransferase, LPCAT
- chlorophyll a/b binding protein, CBP
- cyl-CoA:diacylglycerol acyltransferase, DGAT
- diacylglycerol kinases, DAGK
- Gene Ontology, GO
- Glutathione, GSH
- Jasmonic acid, JA
- jasmonoyl isoleucine, JA-Ile
- lyso-phosphatidylethanolamine acyltransferase, LPEAT
- Osmotic adjustment, OA
- Reactive oxygen species, ROS
- Salicylic acid, SA
- Triacylglycerol, TAG

